# Cryo-EM Structure of the Human Mitochondrial Translocase TIM22 Complex

**DOI:** 10.1101/869289

**Authors:** Liangbo Qi, Qiang Wang, Zeyuan Guan, Yan Wu, Jianbo Cao, Xing Zhang, Chuangye Yan, Ping Yin

**Affiliations:** National Key Laboratory of Crop Genetic Improvement and National Centre of Plant Gene Research, Huazhong Agricultural University, Wuhan 430070, China; Public Laboratory of Electron Microscopy, Huazhong Agricultural University, Wuhan, China; Department of Biophysics, and Department of Pathology of Sir Run Run Shaw Hospital, Zhejiang University School of Medicine, Center of Cryo Electron Microscopy, Zhejiang University, Hangzhou, China; Beijing Advanced Innovation Center for Structural Biology, Tsinghua-Peking Joint Center for Life Sciences, School of Life Sciences, Tsinghua University, Beijing 100084, China

## Abstract

Mitochondria play vital functions in cellular metabolism, homeostasis, and apoptosis^1-3^. Most of the mitochondrial proteins are synthesized as precursors in the cytosol and imported into mitochondria for folding or maturation^4,5^. The translocase TIM22 complex is responsible for the import of multiple hydrophobic carrier proteins that are then folded in the inner membrane of mitochondria^6-8^. In mammalian cells, the TIM22 complex consists of at least six components, Tim22, Tim29, AGK, and three Tim chaperones (Tim9, Tim10a and Tim10b)^9-14^. Here, we report the cryo-EM structure of the human translocase TIM22 complex at an overall resolution of 3.7 angstrom. The core subunit, Tim22, contains four transmembrane helices, forming a partial pore that is open to the lipid bilayer. Tim29 is a single transmembrane protein that provides an N-terminal helix to stabilize Tim22 and a C-terminal intermembrane space (IMS) domain to connect AGK and two TIM chaperone hexamers to maintain complex integrity. One TIM hexamer comprises Tim9 and Tim10a in a 3:3 molar ratio, and the other consists of two Tim9 units, three Tim10a units, and one Tim10b unit. The latter hexamer faces the intramembrane region of Tim22, likely providing the dock to load the precursors to the partial pore of Tim22. Our structure serves as a molecular basis for the mechanistic understanding of TIM22 complex function.

---

Mitochondria are essential eukaryotic cellular organelles with multiple vital functions, such as energetics, metabolism, and cellular signaling^1,2^. These functions are performed by more than 1,000 proteins^3,5^. However, only a small set of these proteins are synthesized in the mitochondria; most of the mitochondria functioning proteins (∼99%) are encoded by nuclear genes, and imported into the correct mitochondrial compartment by specific preprotein translocase complexes^3,5,15,16^. These complexes include the translocase of outer membrane (the TOM complex)^17-20^, the carrier translocase of inner membrane complex (the TIM22 complex)^21-24^, the presequence translocase of the inner membrane (the TIM23 complex)^4,19,25^, the sorting and assembly machinery (the SAM complex)^26,27^, and the mitochondrial import complex (the MIM complex)^28,29^, etc. These translocation machineries are crucial for mitochondrial biogenesis, dynamics, and function^3,4,30,31^. Their dysregulations are connected to dozens of rare mitochondrial diseases and disorders, suggesting an involvement in disease pathogenesis^3,31-33^.

The carrier pathway is one major protein import pathway that is responsible for the translocation and insertion of carrier proteins into the mitochondrial inner membrane^4,5,34-38^. The carrier precursors pass through the pore of the TOM complex, and are then transferred by a hexameric small TIM chaperon complex (the Tim9/Tim10a complex is a main form) to the TIM22 complex^4,5^. Consequently, the TIM22 complex mediates the insertion and lateral release of precursors into the inner membrane in a membrane potential-dependent manner^39^. In addition, the TIM22 complex also transports other multiple-transmembrane-segments containing inner-membrane proteins, including members of the Tim17, Tim22, and Tim23 family^40,41^.

In human, the TIM22 complex consists of at least six components: a hypothetic channel-forming protein Tim22^9,21,24^; three chaperone components (Tim9, Tim10a and Tim10b)^10,42^; newly identified Tim29^11,12^ and acylglycerol kinase (AGK)^13,14^ (Fig. 1a). Tim22 and small chaperones are well conserved from yeast to mammals^9^, but Tim29 and AGK are specific in metazoan^11-14^. AGK participates in lipid biosynthesis^43^, and mutations in *AGK* gene have been associated with Sengers syndrome^44-48^. Mutations in the *TIMM8A* gene (also known as *DDP1* encoding a small chaperone homologue TIM8A) cause deafness dystonia syndrome^49,50^. Recently, mutations in the *TIM22* gene have been identified to cause early-onset mitochondrial myopathy^51^.

**Fig. 1.**
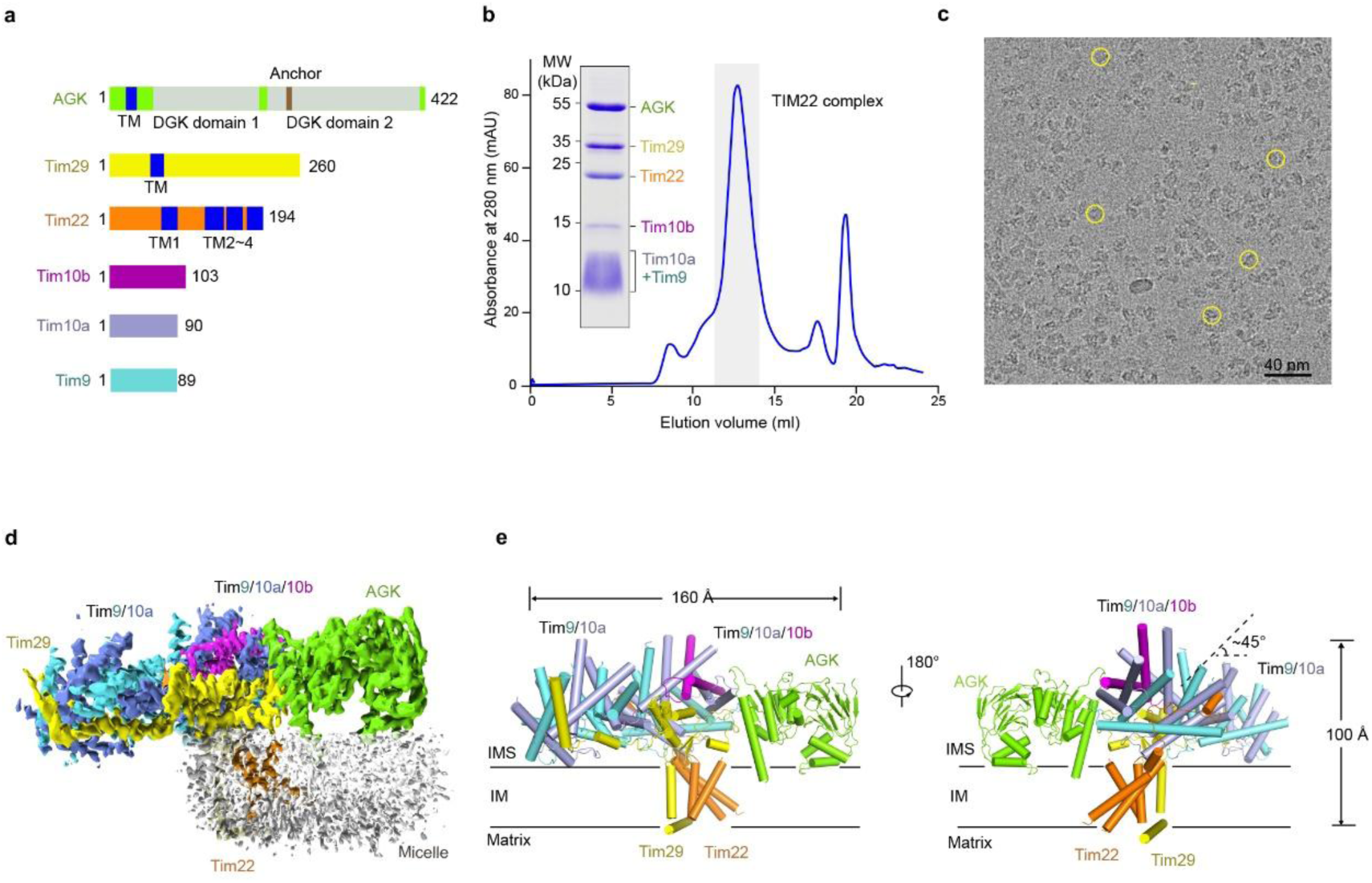
Cryo-EM structure of the human TIM22 complex. **a**, The schematic diagram for each subunit of the TIM22 complex. **b**, A representative gel filtration chromatography of the TIM22 complex. The peak fractions were pooled for cryo-EM study. **c**, A representative electron micrograph of TIM22 complex. The typical particles are marked by yellow circles. **d**, Cryo-EM density map of the TIM22 complex. Tim9, cyan; Tim10a, slate; Tim10b, magenta; Tim22, orange; Tim29, yellow; AGK, chartreuse; the disc-shape micelle, gray. **e**, The opposing side views of the TIM22 complex. The inner membrane (IM) is indicated by two lines between the intermembrane space (IMS) and the matrix. All structure figures were prepared using PyMol.

Despite advances in our understanding of the function and pathophysiology of the TIM22 complex, structural characterization has been sparse. The limited structural information of the TIM22 complex restricted to the crystal structures of Tim9/Tim10a hexameric chaperone and homologues, and a nuclear magnetic resonance (NMR) analysis of carrier precursor associated Tim9/Tim10a^52-55^. Electron microscopy has been also applied to structural studies of translocase complexes. The negatively stained images and the recent cryoelectron microscopy (cryo-EM) structure of the TOM core complex at approximately 6.8 Å revealed pore-forming architecture^56-59^. In contrast, only the low-resolution electron micrographs of the yeast TIM22 complex revealed a two-pore like shape, which was supposed to display a smaller pore diameter than the pores of the TOM core complex^39^.

Here, we report the structure of the human TIM22 complex at resolutions of 3.7 Å for the overall structure and 3.5 Å for the inter-membrane region, determined using single-particle cryo-EM. The structure reveals the assembly and detailed structural information of the complex, and provides an important framework for understanding the function and mechanism of the TIM22 complex in carrier protein maturation.

We co-expressed all six components of the TIM22 complex in human embryonic kidney (HEK) 293F cells. After Flag-tag affinity purification followed by gel filtration, the resulting TIM22 complex displayed good solution (Fig. 1b). The apparent molecular weight was approximately 440 kDa assessed by blue native PAGE (Extended Data Fig. 1), in line to previous findings. Analysis of the purified complex by mass spectrometry (MS) confirmed the presence of all the components of the TIM22 complex. Details of grid preparation, cryo-EM data acquisition, and structural determination of the TIM22 complex can be found in the Methods. Following the initial 2D classification, 3D classification and refinement of the cryo-EM particle images yielded a final 3D EM reconstruction map at an overall 3.7 Å resolution and 3.5 Å resolution for the inter-membrane region of out of 482,959 selected particles (Fig. 1c, Extended Data Fig. 2 and Table 1), according to the gold-standard Fourier shell correlation (FSC) 0.143 criterion (Extended Data Fig. 3). Atomic models were built into the map for Tim22, Tim9, Tim10a, Tim10b, Tim29 and AGK (Fig. 1d and Extended Data Table 2; examples of local densities are shown in Extended Data Fig. 4). The densities for the Tim29 N-terminal helix were of lower resolutions, and poly-Ala was assigned to this region.

The overall structure of the TIM22 complex was approximately 100 Å in height and 160 Å in the longest dimension of width (Fig. 1e). In the structure, there was one Tim22, one Tim29, one AGK, and two hexamer chaperones, Tim9/10a and Tim9/10a/10b, whose stoichiometries were 3:3 and 2:3:1, respectively (Extended Data Fig. 5). Most of the structures were located at the intermembrane space (IMS), including the N-terminus of Tim22 and the large extended C-terminus portion of Tim29 (Extended Data Fig. 5a). Four transmembrane segments (TMs) of Tim22 together with a single TM of Tim29 constitute the transmembrane element at the center of the TIM22 complex. Only one N-terminal helix of Tim29 protrudes from the core transmembrane region and was oriented nearly parallel to the plane of the membrane at the matrix side (Fig. 1e). The Tim9/10a/10b hexamer, similar to a hub, was located at the center of TIM22 complex, and was encircled by the N-terminus of Tim22, the middle portion of Tim29, AGK, and the Tim9/10a chaperone. Interestingly, the Tim9/10a/10b hexamer was not perpendicular to the membrane, but rather approximately 45° tilted (Fig. 1e). The Tim9/10a chaperone and AGK were located at the nearly-opposite side of the Tim9/10a/10b hexamer, and AGK was anchored to the membrane via its portion of the N-terminal helix and an additional membrane anchor helix loop helix (Fig. 1e).

Tim22 contains two helices (α1 and α2) connected by an extended loop 1, and 4 transmembrane segments (TM 1-4) (Fig. 2a and Extended Data Fig. 5a). The 23 N-terminal residues and the matrix loop (residues 94-118; connection between TM1 and TM2) had no EM density, most likely due to their intrinsic flexibility. Two helices (α1 and α2) protrude toward the intermembrane space and interact with the Tim9/10a/10b hexamer. Helix α1 and loop 1, similar to a hook, sinuously wind around the groove between the inner helices and the outer helices of Tim9 and Tim10a. Helix α1 was nestled in a greasy pocket formed by Tim9 and Tim10a (Fig. 2b), and likely functioned as a plug to obstruct the hydrophobic carrier precursor from wedging within the hexamer chaperone from this side (Extended Data Fig. 6a). The recently identified disease-related mutation (Val33Leu) of Tim22 was located in the helix α1^51^.

**Fig. 2.**
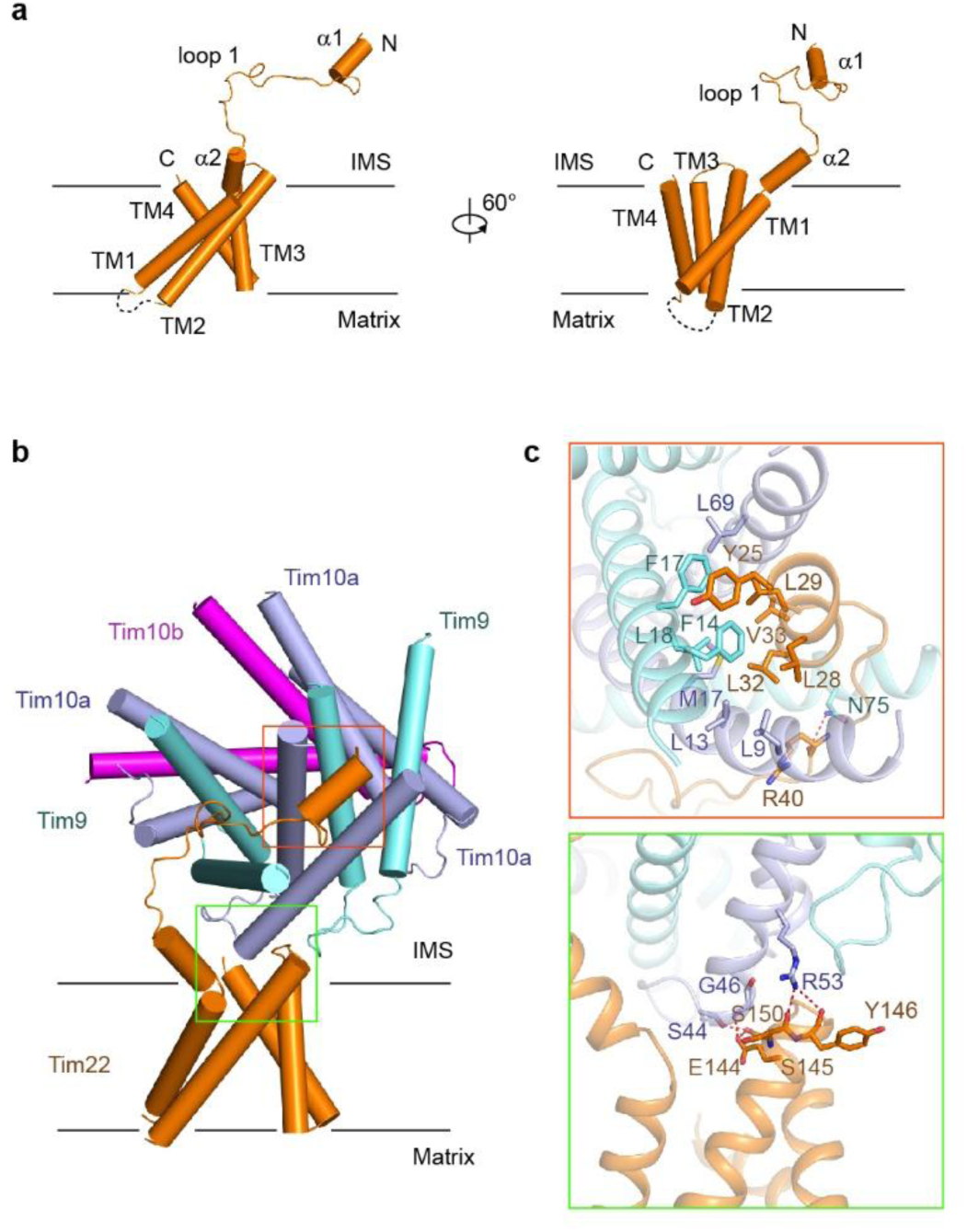
Structure of the Tim22 subunit. **a**, Two side views of Tim22. The transmembrane (TM) segments are labeled. **b**, Tim22 contacts the Tim9/10a/10b chaperone mainly via the N-terminal helix in the intermembrane space and the helices bundle at the inner membrane. The two interfaces are highlighted by red and green box. **c**, Key residues mediating the interactions between Tim22 subunit and Tim9/10a/10b chaperone are shown as sticks.

Specific interactions comprise van der Waals contacts and one hydrogen bond. Hydrophobic residues, Leu28, Leu29, Leu32, and Val33 interacted with residues Leu9, Leu13, Met17, and Leu69 from helices of Tim9, and residues Phe14, Phe17, and Leu18 from the inner helix of Tim10a. The aromatic ring of Tyr25 from Tim22 stacked against Phe14 and Phe17 from Tim10a. Asn75 from Tim10a, was found to form a hydrogen bond with the main chain of Arg40 from Tim22 (Fig. 2c). Reinforcing these interactions, one residue from helix α2 and four residues from the last turn of TM2 of Tim22 contacted residues of the outer helix of Tim9. Glu67 from Tim22 interacted with Lys45 from Tim9 to form a salt bridge. The side chains of Glu144 and Ser150 from Tim22 formed hydrogen bonds with Ser44 and the main chain of Gly46 from Tim9. In addition, the main chains of Ser145 and Tyr146 from Tim22 accepted hydrogen bonds from Arg53 of Tim9 (Fig. 2c).

Notably, only one Tim22 was observed in the complex. The four TMs of Tim22 were unlike a closed pore channel but constituted a lateral hydrophobic cave that was exposed to the lipid bilayer (Fig. 2a), which is reminiscent of insertase^60^. A disulfide bond formed between Cys69 and Cys141 appeared to stabilize the conformations of TM1 and TM2 (Extended Data Fig. 6b, c), which was consistent with biochemical studies. Tim22 homologues from yeast to human share more than 40% similarity, and most of the interacting residues and two cysteines are conserved (Extended Data Fig. 6c), suggesting that the Tim22 homologues exhibit similar folds^9^.

Tim29 was recently identified as a metazoan-specific subunit of the human TIM22 complex, which is required for the stability of the complex. Our structure corroborated this observation, since. Tim29 exhibited an extended conformation comprising of a long N-terminal helix α1 in the matrix, a single TM, an intermembrane space domain (IMS Domain) and a C-terminal chaperone Tim9/10a recruiting motif (CRM) (Fig. 3a and Extended Data Fig. 5a). One phospholipid, which was most likely to be phosphatidylethanolamine, was observed at the IMS (Extended Data Fig. 7a, b). The helix α1 was oriented nearly parallel to the plane of the membrane in the matrix and perpendicular to the TM of Tim29 (Fig. 3a, b), which appeared to interact with TM3 of Tim22 and stabilize it. However, the lack of EM density for the side chains of helix α1 prevented an unambiguous assessment of the details of these molecular interactions. Interestingly, the single TM of Tim29 was positioned far from the TMs of Tim22, of some extent, suggesting this TM was unable to strictly associate with TMs of Tim22 (Fig. 3b). This arrangement suggests the TM is flexible for regulation of precursor insertion.

**Fig. 3.**
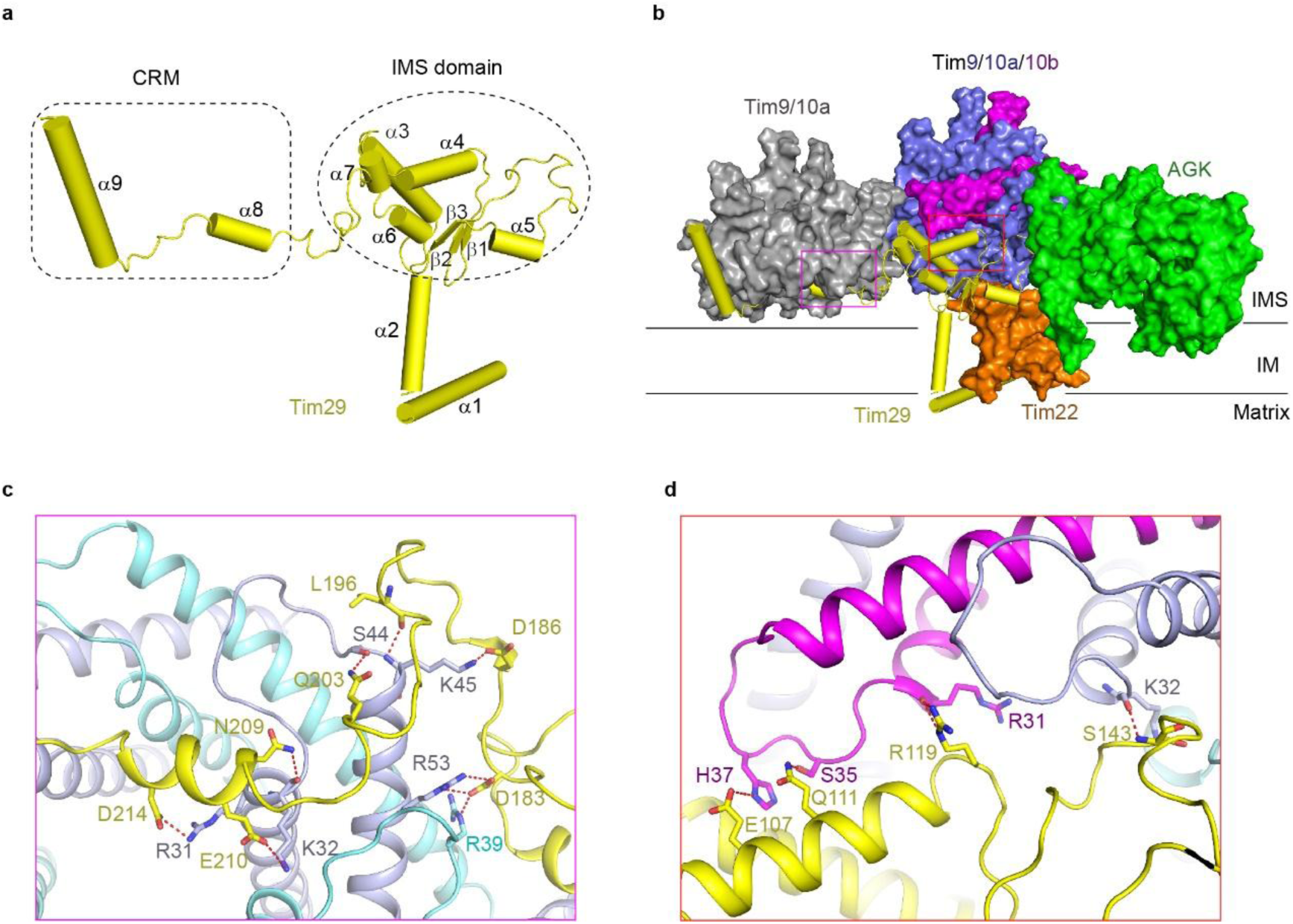
Structure of the Tim29 subunit. **a**, Tim29 subunit forms extensive contacts with both Tim9/10a and Tim9/10a/10b chaperones via the C-terminal segments. The interacted regions are indicated by pink and red box, respectively. **b**, The overall structure of the Tim29 subunit. The C-terminus of Tim29 is divided into two segments: IMS (intermembrane space) domain and CRM (C-terminal recruiting motif). The secondary structural elements are labeled. **c, d**, Key residues mediating the interactions between Tim29 subunit and Tim9/10a (**c**) or Tim9/10a/10b (**d**) are shown as sticks.

The IMS domain of Tim29 interacts with the Tim9/10a/10b hexamer mainly through the helix α4, as illustrated by three polar residues Glu107, Gln111, and Arg119 hydrogen-bonding with His37, Ser35, Arg31 from Tim10b, respectively (Fig. 3b, d). Additionally, the main chains of Ser143 from Tim29 and Lys32 from Tim10a form a hydrogen bond (Fig. 3d). These interactions robustly fix the orientation of the Tim9/10a/10b hexamer, in part due to the presence of only one Tim10b in the heterohexamer (Fig. 3b and Extended Data Fig. 5a).

The CRM of Tim29 makes contact with the Tim9/10a chaperone via extensive electrostatic interactions (Fig. 3b). The negatively charged residues of Asp183, Asp186, Glu210 and Asp214 interact with several positively charged residues, Arg39 from Tim9, including Arg53, Lys 45, Lys32, and Arg31 from Tim10a, respectively (Fig. 3c). Interestingly, the external C-terminal helix, α9, has no interaction with the Tim9/10a chaperone. Given that this portion is rich in negatively charged residues (Extended Data Fig. 7c, d), it might recruit the other Tim9/Tim10a chaperone or interact with the TOM complex, which has been shown to occur biochemically^12^.

The Tim9/10a chaperone and the Tim9/10a/10b chaperone share almost identical symmetric hexamer structures as that of the reported free Tim9/10a hexamer^52^ (Fig. 4 and Extended Data Fig. 8a). Each individual subunit displays a helix-loop-helix fold and is stabilized by two intramolecular disulfide bonds from a highly conserved “twin CX_3_C motif” (Fig. 4b). The six inner helices (helix α1) collectively form a donut-shape with a 14∼16 Å diameter central hole. The outer helices (helix α2) act as tentacles that radiate from the central region (Fig. 4a. b). Superimposing Tim9, Tim10a, and Tim10b revealed that the less conserved connecting loops (C loops) exhibit distinct conformations (Fig. 4c), in line to the observation that the C loop of Tim10b is primarily responsible for interaction with Tim29 (Fig. 3d). Furthermore, the C-terminus of the outer helix of Tim10b tilts towards the inner helix (Fig. 4b), mainly due to its Pro72 residue (Fig. 4d). Thus, the major conformational difference between these two chaperones appears to be the outer helix of Tim10b, while the diameter of the Tim9/10a hole appears to be subtly narrowed (Extended Data Fig. 8b). A circular groove was observed between the inner and outer helices, which was previously hypothesized to hold unstructured carrier precursors^52,55^. The twist outer helix of Tim10b disrupts the continuous groove (Extended Data Fig. 8c, d), which might be involved in precursors unloading from the Tim9/10a chaperone. The Tim9/10a chaperone and the Tim9/10a/10b hexamer were selectively bound to the CRM and IMS domain of Tim29 as determined by Tim10b (Fig. 3c, d). Tim10b interacted with the IMS domain of Tim29 and the DGK domain of AGK (described below). These specific interactions prevented the binding of the Tim9/10a chaperone in this position (Fig 3b).

**Fig. 4.**
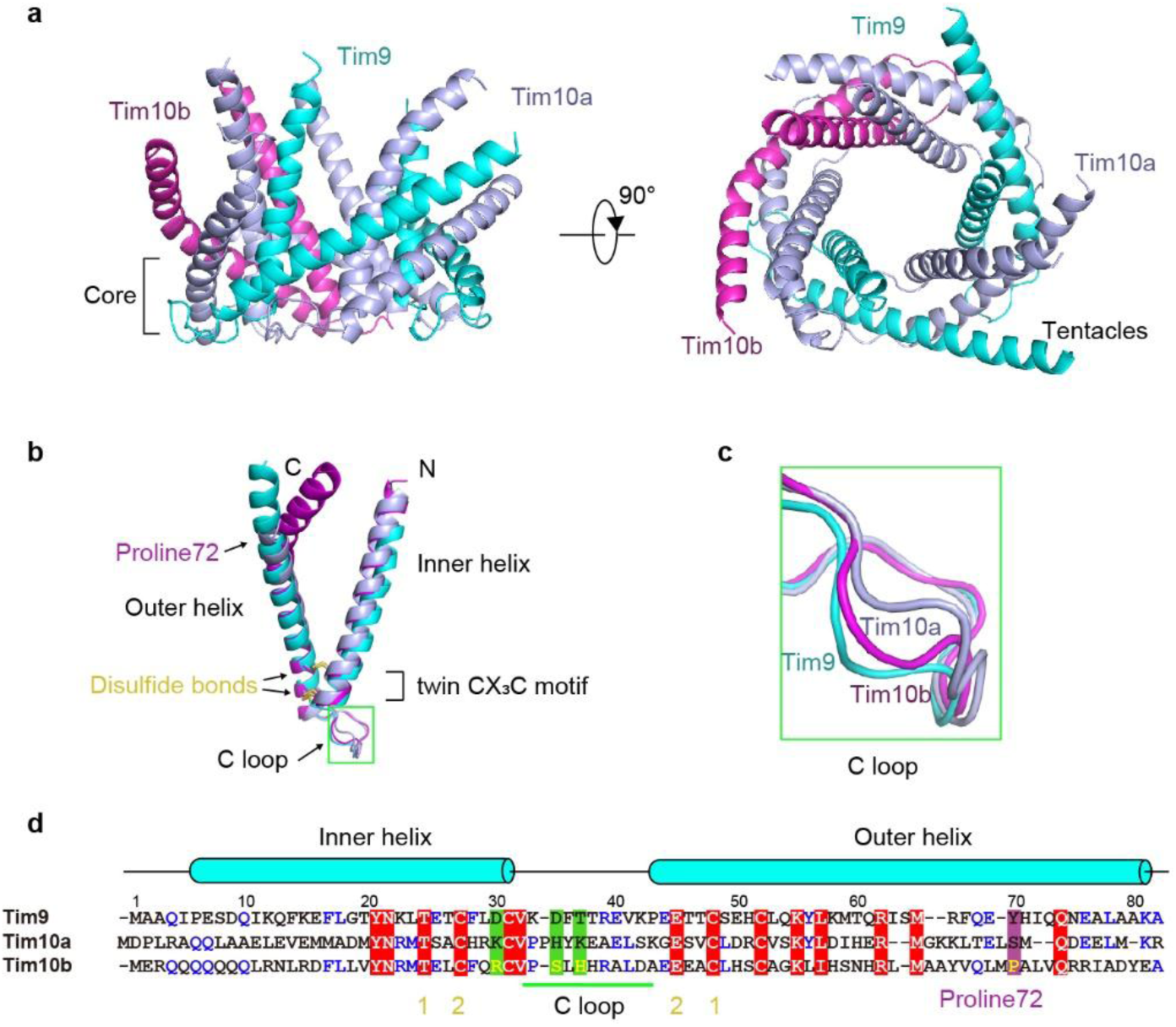
Structure of the Tim9/10a/10b hexamer. **a**, Two perpendicular views of Tim9/10a/10b hexamer. The Tim9/10a/10b hexamer forms a pore. **b**, Structural superimposition of Tim9, Tim10a and Tim10b. Disulfide bonds formed from the signature cysteines of twin CX_3_C motif are indicated. The C (center) loop is highlighted by green box. **c**, Structural alignment of the C-loop of Tim9, Tim10a, and Tim10b. **d**, Sequence alignment of Tim9, Tim10a and Tim10b. The conserved residues are colored red. Residues in the C loop involved in the interaction with the Tim29 subunit are colored green. Cysteines in the twin CX_3_C motif forming the disulfide bonds are shown with yellow numbers. Proline72 is colored purple. The inner and outer helices are drawn above the sequences.

AGK is a mitochondrial lipid kinase that converts monoacylglycerol (MAG) and diacylglycerol (DAG) to lysophosphatidic acid (lyso-PA) and phosphatidic acid (PA), respectively^43^. Similar to the DAG kinase homologue, DgkB from *Staphylococcus aureus* (SaDgkB), the AGK structure exhibits a typical two-domain fold^61^ (Fig. 5a and Extended Data Fig. 5b). Notably, a predicted TM (helix α1) preceding domain 1 is partially embedded in the membrane (Fig. 5a), and is consistent with previous findings^13,14^. A protruded helix α9 and an ensuing loop of domain 2 were found to be anchored to the membrane via Trp225, Tyr226, L227, L230, Phe237 and Phe238 (Fig. 5a), generating a positively-charged cavity between AGK and the membrane to potentially facilitate the carrier precursor insertion (Fig. 5b). These membrane-anchoring structural features were found to be AGK-specific (Fig. 5c). AGK interacts with both Tim29 and the Tim9/10a/10b hexamer, corroborating previous studies that suggested AGK is a *bona fide* subunit of the TIM22 complex. Arg40 from helix α1 offers hydrogen bonds to the main chains of Tyr151 and Gln153 from Tim29 (Fig. 5d). Gln52, Ala58, and Asp94 were found to hydrogen-bond with Lys45 from Tim10a, Arg62 from Tim10b, and Arg39 from Tim9, respectively (Fig. 5e).

**Fig. 5.**
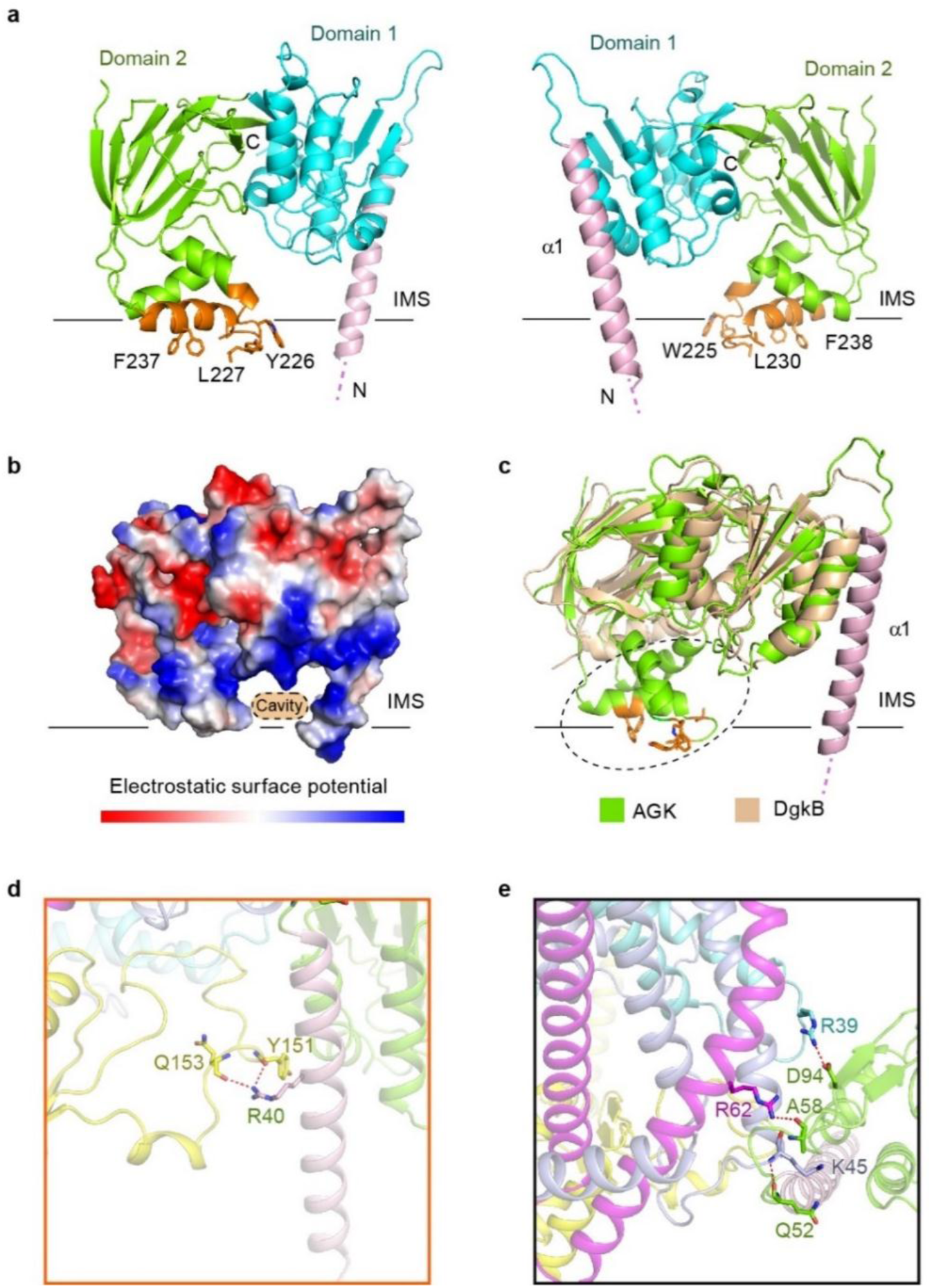
Structure of the AGK subunit. **a**, Two opposing views of AGK, which is composed by α1, domain 1 and domain 2. The first α-helix (light pink) is embedded into the membrane. Key residues bound to the membrane are shown as orange sticks. **b**, A positively charged cavity of AGK is formed in between the first α-helix and the anchor. **c**, Structural alignment of AGK (*Homo sapiens*) and DgkB (*Staphylococcus aureus*, PDB code: 2VQ7). AGK exhibits additional membrane anchor (a dashed circle) and the transmembrane α1. **d, e**, Key residues involved in the interactions between AGK and Tim29 (**d**) or Tim9/10a/10b chaperone (**e**) are shown as sticks.

The structure of the TIM22 complex presented here not only offers the first molecular level view of the interactions that facilitate the mammalian TIM22 complex assembly, but also provides a framework for a mechanistic understanding of the import of carrier proteins through the TIM22 complex. Typically, the carrier protein contains six TMs that are assembled in three repeats, each comprising two TMs connected by a short matrix helix^62-65^. Both the amino and carboxy termini are oriented towards the intermembrane space^63,64^. And, the precursor protein is wrapped into the hydrophobic groove of the Tim9/10a chaperone^55^. Based on our structural analysis, we proposed a “bending-in” insertion model, wherein Tim22 and AGK generate a space-limited environment to recruit the two transmembrane helices of the carrier proteins at once, and then bend them into the membrane (Extended Data Fig. 9). After insertion of one repeat into the membrane, the two TM segments intermediate might be transiently stabilized by the Tim22 until the next repeat inserts. This scenario the structures of the TIM22 complex with the carrier precursor during an insertion cycle are necessitated to fully understand the carrier translocation mechanism.

## Data availability

Cryo-EM map for the human TIM22 complex is available on the Electron Microscopy Data Bank under accession number EMD-9958. Coordinate of the atomic structure have been deposited in the Protein Data Bank under accession number 6KC9. All other data are available from the corresponding authors upon reasonable request.

## Methods

### Transient expression of the human TIM22 complex

The codon-optimized full-length cDNAs for subunits of the human TIM22 complex were synthesized by General Biosystems Company (Tim22, Uniport: Q9Y584; Tim29, Uniport: Q9BSF4; Tim9, Uniport: Q9Y5J7; Tim10a, Uniport: P62072, Tim10b, Uniport: Q9Y5J6; AGK, Uniport: Q53H12). TIM22, TIM29 and AGK were subcloned into the plasmid A; TIM9, TIM10A and TIM10B were subcloned into the plasmid B using the pMlink vector^1^. Tim22 contains a N-terminal triple Flag tag. Expi293F^TM^ (Invitrogen) cells were cultured in SMM 293TI medium (Sino Biological Inc.) at 37 °C under 5% CO _2_ in a shaker and diluted into 2.0 × 10^6^ cells ml^-1^ with fresh medium when the cell density reached 3.5 × 10^6^ ∼ 4.0 × 10^6^ cells ml^-1^ for further transfection. For 1 liter cell culture, 1.4 mg plasmid A and 0.6 mg plasmid B were pre-incubated with 4 mg linear polyethylenimines (PEIs) (Polysciences) in 50 ml fresh medium for 20 min. The transfection was initiated by adding the mixture into the diluted cell culture. Transfected cells were cultured for 48 hours before harvesting.

### Mitochondria Preparation

The cells were harvested by centrifugation at 800 g for 20 min and washed with PBS, then resuspended in the buffer A containing 10 mM Tris-HCl pH 7.5, 70 mM sucrose, 210 mM mannitol, 1 mM EDTA, 1mg ml^-1^ BSA and 1mM PMSF^2^. The cells were disrupted using dounce homogenizer (sigma) for 80 cycles on ice and the homogenate was centrifuged at 3,000 g for 10 min. The supernatant was further centrifuged at 20,000 g for 20 min to obtain the crude mitochondrial pellet. The pellet was resuspended in buffer B containing10 mM Tris pH 7.5, 250 mM sucrose, 60 mM KCl and 0.1 mM EDTA and centrifuged at 60,000 g at 4 °C for 20 min in a discontinuous Percoll density gradient^3^. The clear mitochondria layer was obtained carefully and diluted with buffer C containing 10 mM Tris-HCl pH 7.5, 70 mM sucrose, 210 mM mannitol. The highly pure mitochondria were collected by centrifugation at 20,000 g for 30 min and stored in buffer C at −80 °C before use.

### Purification of the TIM22 complex

The TIM22 complex were extracted from pure mitochondria by 1% LMNG (Anatrace), 0.25% soybean lipids (Sigma) and 0.1 % CHS (Anatrace) in lysis buffer containing 25 mM HEPES pH 7.4, 100 mM KOAc, 10 mM Mg(OAc)_2_, 0.1 mM EDTA, 10% glycerol, 1 mM PMSF, 4 µg ml^-1^ pepstatin A, 4 µg ml^-1^ aprotinin and 5 µg ml^-1^ leupeptin at 4 °C for 1.5 h^4^. The extraction was centrifuged at 100,000 g for 30 min to remove the unsoluble component. The supernatant was incubated with anti-Flag G1 affinity resin (Genscript) at 4 °C for 2 h and then washed with 30 bed volumes of lysis buffer added 0.05% GDN (Anatrace). The protein was eluted with lysis buffer added 0.05% GDN and 300 μg ml^-1^ Flag peptide (Genscript). The protein solution was concentrated with a 100-kDa cut-off centricon (Milipore) and further purified by Superose-6 increase 10/300 column (GE Healthcare) using a buffer containing 50 mM Imidazole pH 6.0, 150 mM NaCl, 2 mM MgCl_2_ and 0.05% GDN^5^. The peak fractions were pooled and concentrated to 7 mg ml^-1^ for further cryo-EM study.

### Blue native-PAGE analysis

Blue native PAGE technique was used to determine native TIM22 complex mass as described previously^6^. Chromatographically purified TIM22 complex sample was mixed with 10 × loading buffer (0.1% (w/v) Ponceau S, 50% (w/v) glycerol) and subjected to 4%-16% blue native PAGE mini gel (Invitrogen) for electrophoresis at 4 °C. The TIM22 complex was transferred onto a PVDF membrane and detected by immunoblotting using Tim22 antibody (cat number: 14927-1-AP, Proteintech). The size of the TIM22 complex was determined based on the mobility pattern of a native protein molecular weight standard (cat number: LC0725, Invitrogen).

### Mass spectrometry analysis

The TIM22 complex proteins were separated by 1D SDS-PAGE, the gel bands of interest were excised from the gel, reduced with 5 mM of dithiotreitol, and alkylated with 11 mM iodoacetamide. In gel digestion was then carried out with sequencing grade modified trypsin in 50 mM ammonium bicarbonate at 37 °C overnight. The peptides were extracted twice with 0.1% trifluoroacetic acid in 50% acetonitrile aqueous solution for 30 min. Extracts were then centrifuged in a speedvac to reduce the volume. Tryptic peptides were redissolved in 20 μl 0.1% TFA and analyzed by LC-MS/MS.

### Cryo-EM Data Acquisition

For cryo-EM sample preparation, 3.5 μl aliquots of the recombinant human TIM22 complex (7 mg ml^-1^) were dropped onto glow discharged holey carbon grids (Quantifoil Au R1.2/1.3, 300 mesh), blotted with a Vitrobot Mark IV (ThemoFisher Scientific) using 5 s blotting time with 100% humidity at 8 °C, and plunged into liquid ethane cooled by liquid nitrogen. The sample was imaged on an FEI Titan Krios transmission electron microscope at 300 kV with a magnification of 29,000 ×. Images were recorded by a Gatan K2 Summit direct electron detector using the counting mode. Defocus values varied from −1.8 to −2.5 μm. Each image was dose fractionated to 32 frames with a total electron dose of 60 e^-^ Å^-2^ and a total exposure time of 8.0 s. SerialEM^7^ was used for fully automated data collection. All stacks were motion corrected using MotionCor2^8^ with a binning factor of 1, resulting in a pixel size of 1.014 Å and dose weighting was performed concurrently. The defocus values were estimated using Gctf^9^.

### Preliminary data processing

1,951 micrographs were used for initial data analysis. A small sets of particles for the TIM22 complex were auto-picked using reference-free autopicking method with Laplacian-of-Gaussian in RELION3.0^10^. Good 2D averages were generated after 2D classification and 552,779 particles were auto-picked with selected good 2D averages as template. One round of 2D classification was performed, remaining 305,689 good particles. Initial model was generated with stochastic gradient descent method in RELION^10^ using about 1000 particles. Global 3D classification was performed with this initial model as reference, generating one good class with 36.8% of input particles and three bad classes.

### Data processing of the TIM22 complex

For the complete data set of the TIM22 complex, 2,401,464 particles were auto-picked from 14,105 micrographs. A guided multi-reference global classification procedure was applied to the full dataset. One good and five bad references from preliminary data processing were used as the initial references (Round 1). These six references were low-pass filtered to 40 Å. To avoid the problem of discarding good particles, we simultaneously performed three parallel multi-reference 3D global classifications. After the global classification, particles that belong to the good classes from 24/27/30 iterations of the three parallel runs were separately used as the input for follow-up local classification. Six 3D models from global classification were used as references for the multi-reference local classification. Good particles from these 3×3 parallel runs were merged, and the duplicated particles were removed as described previously^11^. 1,107,389 particles (46.1% of the original input) remained for following processing.

A second round (Round 2) of multi-reference local 3D classification was performed with five references (from global classification, scale to pixel size 2.028 Å) using re-extracted and re-centered 2x binned particles (pixel size: 2.028 Å). Particles from the good classes (representing 45.4/47.8/50.2% of the input particles) were combined to yield 621,049 particles (representing 25.9% of the total original particles).

A third round (Round 3) local classification was performed using unbinned particles (pixel size: 1.014 Å), focusing on the inter-membrane region. The largest class of 482,959 particles (representing 77.8% of the input particles or 20.1% of the total original particles) yielded an average resolution of 3.73 Å after auto-refinement. With application of a soft mask on the inter-membrane region, these particles gave an average resolution of 3.53 Å for this region after continuous auto-refinement.

In the 3.73Å overall and 3.53Å intermembrane region maps of the TIM22 complex, the local resolution reaches 3.2-3.5 Å in the core regions. The angular distributions of the particles used for the final reconstruction of the TIM22 complexes are reasonable, and the refinement of the atomic coordinates did not suffer from severe over-fitting. The resulting EM density maps display distinguishing features for the amino acid side chains in most regions.

Reported resolutions were calculated on the basis of the FSC 0.143 criterion, and the FSC curves were corrected for the effects of a soft mask on the FSC curve using high-resolution noise substitution^12^. Prior to visualization, all density maps were corrected for the modulation transfer function (MTF) of the detector, and then sharpened by applying a negative B-factor that was estimated using automated procedures^13^. Local resolution variations were estimated using RELION^10^.

### Model building and refinement

We combined homology modeling and de-novo model building to generate the atomic models.

Identification and docking of the two chaperones Tim9/10a and Tim9/10a/10b were facilitated by the crystal structure of the human mitochondrial Tim9/10a hexametric complex (PDB code: 2BSK). Tim10b was manually identified by comparing the sequence variance of these three proteins and the cryo-EM density, and then manually build by COOT^14^.

Identification of AGK protein was facilitated by the homology structure of a diacylglycerol kinase DgkB from *Staphylococcus aureus* (PDB code: 2QV7). The atomic model of AGK were generated by CHAINSAW^15^ and the backbone was manually adjusted using COOT^14^. After that, automated model rebuilding was performed with RosettaCM using the adjusted model as the template and the experimental cryo-EM density as a guide^16-18^. Then the hydrogen atoms of the generated model were removed and model building was further performed manually using COOT^14^, The single trans-membrane helix and a membrane anchoring motif was de-novo built using COOT^14^.

The intermembrane space (IMS) domain of Tim29 was located under the Tim9/10a/10b hexamer. The local resolution of IMS domain in cryo-EM map reaches around 3.2 Å, allowing us de-novo building of these regions. The transmembrane helix and a C-terminal fragment which recruits the Tim9/10a chaperone was also de-novo built by COOT^14^. A N-terminal helix of Tim29 was identified to be located in the matrix, adhering to the membrane and stabilizing the four transmembrane helices Tim22. Due to limited resolution (5∼7 Å), only poly-ALA helix was built for the N-helix.

Tim22 is supposed to have four transmembrane helices^19^. The cryo-EM density of the membrane region is relatively lower than the soluble region, ranging from 5 Å to 3.2 Å. Luckily, the local resolution of the loop between TM2 and TM3 is high enough, around 3.5 Å, which allow us identify three bulky residues (Y146, R147, and W152) in the linker. The density of the loop connecting TM3 and TM4 can also be clear visualized, which contacts with N-helix of Tim29 in the mitochondrial matrix. A potential S-S bond between TM1 and TM2 can be clearly visualized, which help us assign the sequence. A N-terminal plug bound to Tim9/10a/10b hexamer was also manually built by COOT^14^.

The final model of the TIM22 complex was refined against the overall cryo-EM maps using PHENIX^20^ in real space with secondary structure restraints. Overfitting of the overall model was monitored by refining the model in one of the two independent maps from the gold-standard refinement approach, and testing the refined model against the other map^21^. The structures of the TIM22 complex were validated through examination of the Molprobity scores and statistics of the Ramachandran plots. Molprobity scores were calculated as described^22^.

## ACKNOWLEDGMENTS

We thank Dr. Chang at Center of Cryo Electron Microscopy, Zhejiang University, for assistance during data collection; and research associates at the Center for Protein Research and Public Laboratory of Electron Microscopy, Huazhong Agricultural University, for technical support. We thank the Tsinghua University Branch of China National Center for Protein Sciences (Beijing) for providing the technical support on the Cryo-EM and High-Performance Computation platforms. We thank Meng Han and protein chemistry Facility at the Center for biomedical Analysis of Tsinghua University for sample mass-spec analysis. We thank Ms. Wang Yan for model figure design. This work was supported by funds from the Ministry of Science and Technology of China (2018YFA0507700), the National Natural Science Foundation of China (31722017), the Fok Ying-Tong Education Foundation (151021), and the Fundamental Research Funds for the Central Universities (2662017PY031). This research was supported by Beijing Advanced Innovation Center for Structural Biology (to Dr. Chuangye Yan).

## AUTHOR CONTRIBUTIONS

P.Y. conceived the project. L.Q., Q.W., P.Y. designed all experiments. L.Q., Q.W., and Y.W. performed the experiments. Q.W., Z.G., J.C., and X. Z collected the EM data. Z.G. and C. Y. determined the structure. All authors analyzed the data and contributed to manuscript preparation. P.Y wrote the manuscript.

---

**Extended Data Fig. 1.**
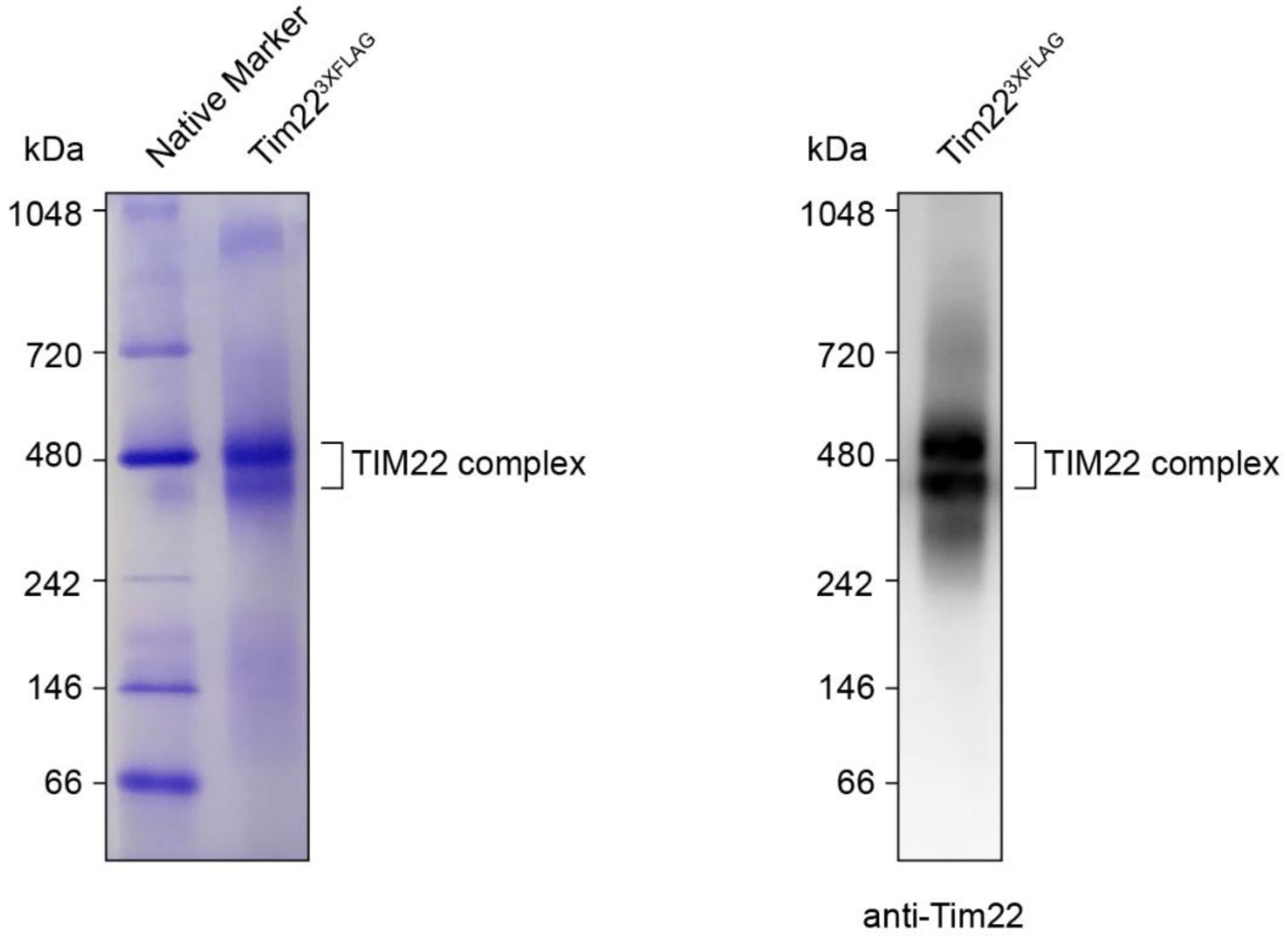
Blue native PAGE analysis of the TIM22 complex. Purified TIM22 complex sample was analyzed by Blue native PAGE. The complex bands were visualized by Coomassie blue staining (left panel) and western blot (right panel).

**Extended Data Fig. 2.**
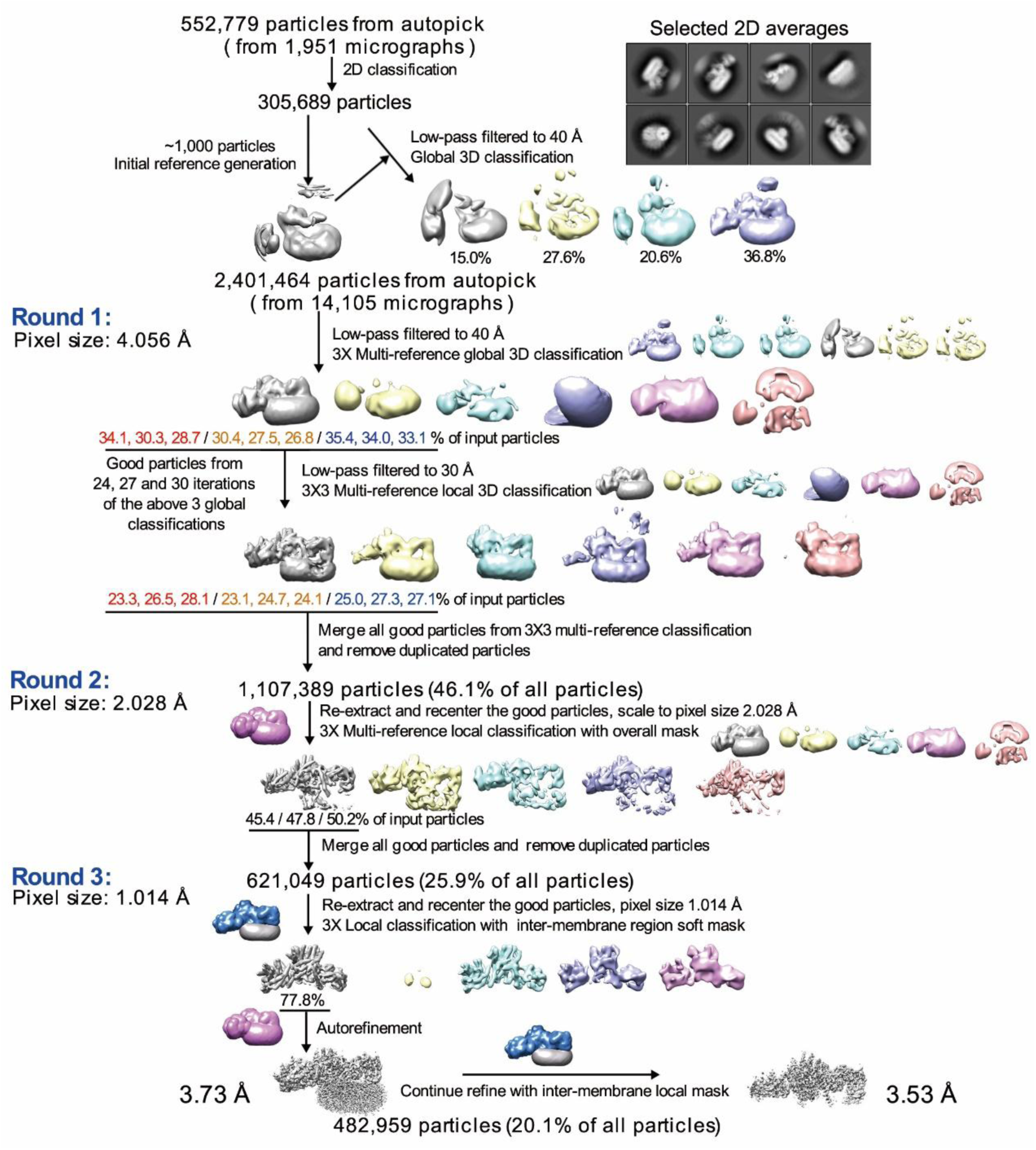
Flowchart for cryo-EM data processing of the human TIM22 complex. On the basis of the FSC value of 0.143, the final reconstruction has an average resolution of 3.73 Å for the overall map, 3.53 Å for the inter-membrane region. Details are presented in Methods.

**Extended Data Fig. 3.**
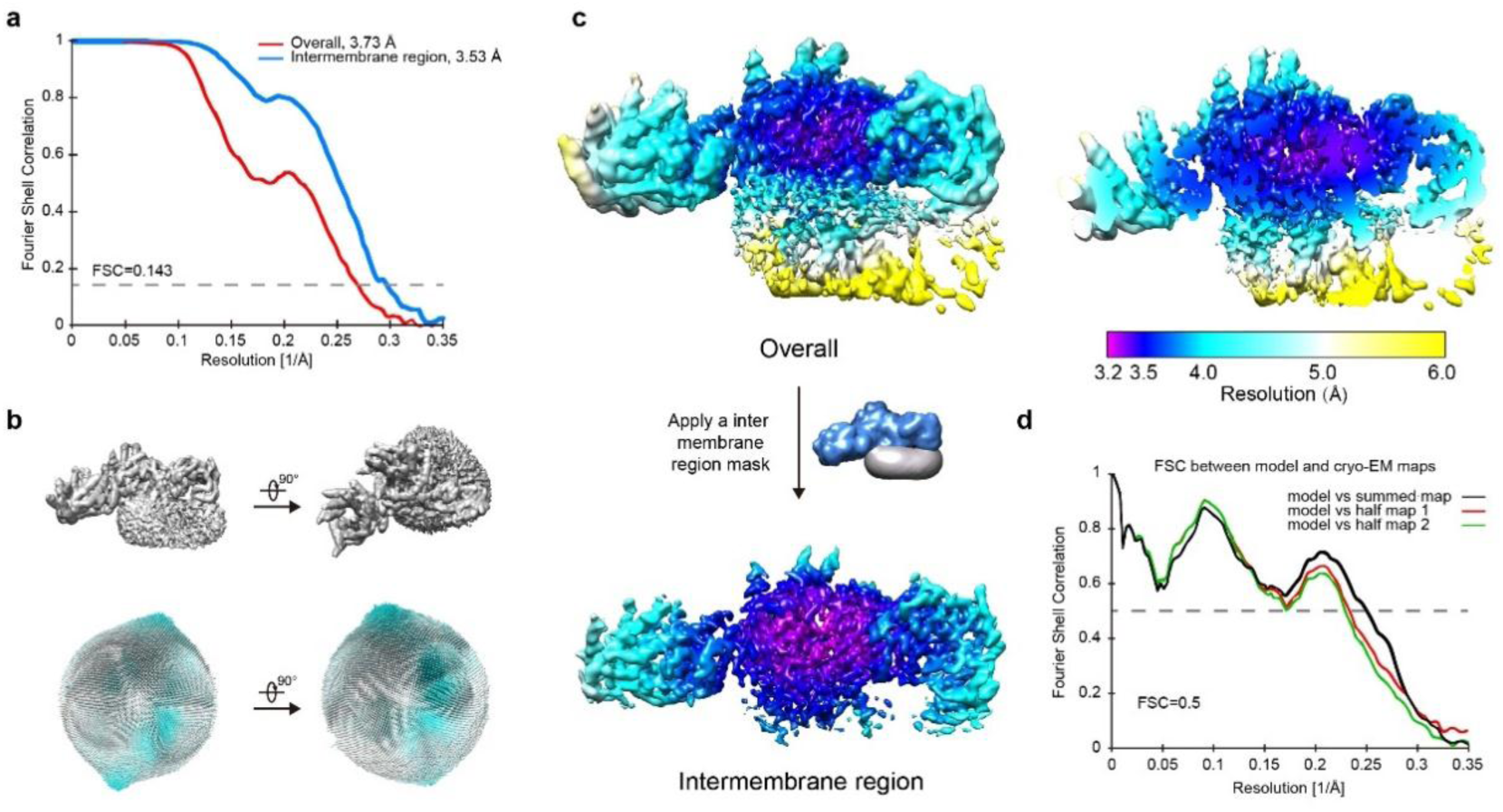
Cryo-EM analysis of the human TIM22 complex. **a**, The average resolutions for the TIM22 complex are estimated to be 3.73 Å and 3.53 Å for the overall and inter-membrane region, respectively. The resolutions are reported on the basis of the FSC criterion of 0.143. **b**, Angular distribution of the particles used for reconstruction of the TIM22 complex. Each cylinder represents one view and the height of the cylinder is proportional to the number of particles for that view. **c**, The local resolutions are color-coded for the TIM22 complex (left panel) and the inter-membrane region (right upper panel). The highest resolution of the EM maps reaches 3.2 Å. **d**, The FSC curves of the final refined models of the TIM22 complex versus the overall maps that it is refined against (black); of the model refined in the first of the two independent maps used for the gold-standard FSC versus that same map (red/purple); and of the model refined in the first of the two independent maps versus the second independent map (green/blue). The generally similar appearances between the purple and blue, red and green curves indicate that the refinement of the atomic coordinates did not suffer from severe over-fitting.

**Extended Data Fig. 4.**
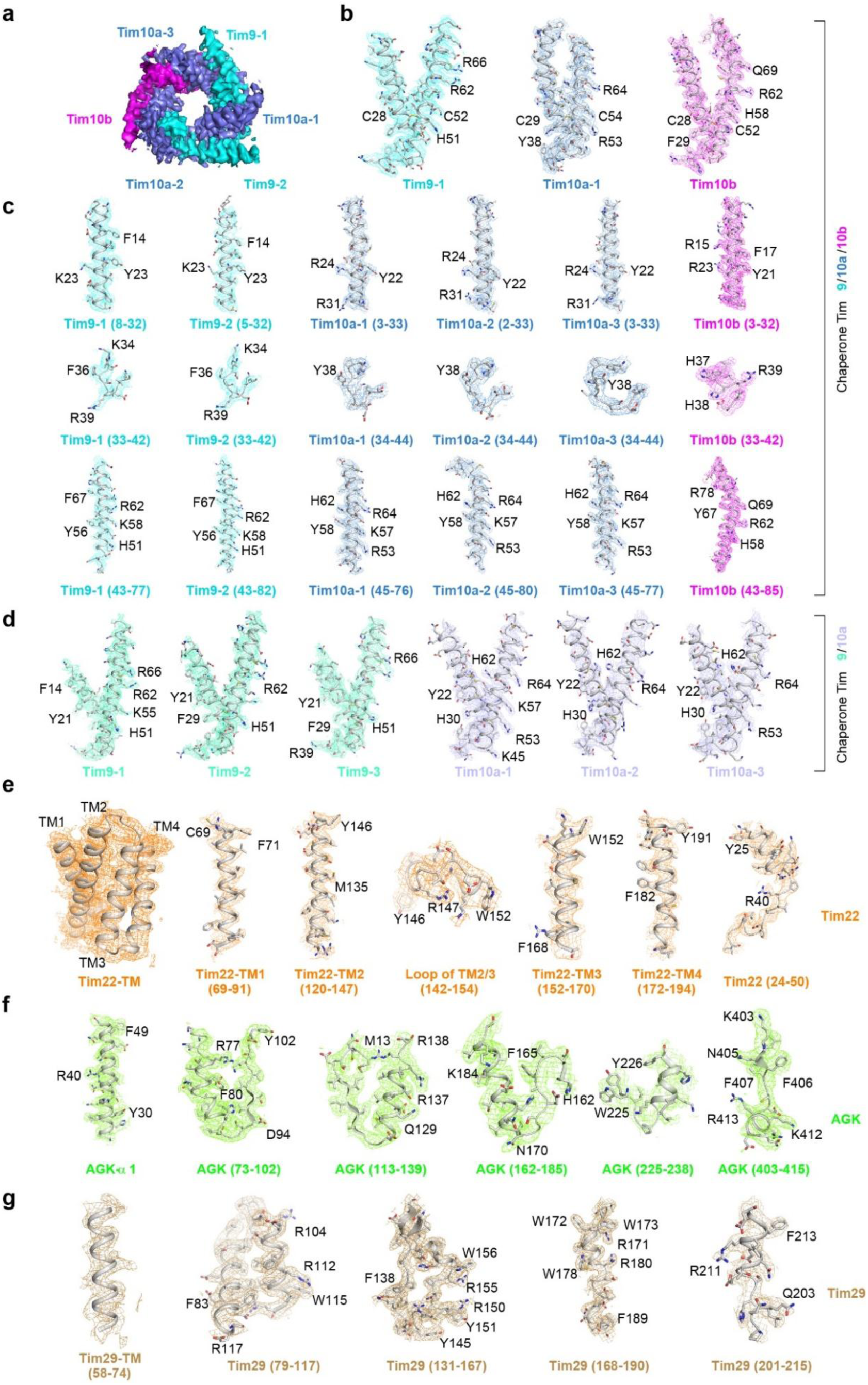
The electron microscopy maps for the subunits of the TIM22 complex. **a**-**c**, The electron microscopy maps for the Tim9/10a/10b hexamer: Overview of chaperone Tim9/10a/10b, composed of Tim9, Tim10a and Tim10b in a ratio of 2:3:1 (**a**); A close up view for representative Tim9, Tim10a and Tim10b (**b**); Three segments (helix-loop-helix) of Tim9, Tim10a and Tim10b (**c**). **d**, The electron microscopy maps for chaperone Tim9/10a. **e**, The 4 transmembrane segments (TM1-4), intermembrane space segments (24-50) and loop (142-154) of Tim22. **f**, AGK α-1 (the predicted TM), 225-238 (the membrane-anchored region), 73-102, 113-139, 162-185, and 403-415 of AGK. **g**, The transmembrane segments (58-74) and intermembrane space segments (79-117, 131-167, 168-190, 201-215) of Tim29.

**Extended Data Fig. 5.**
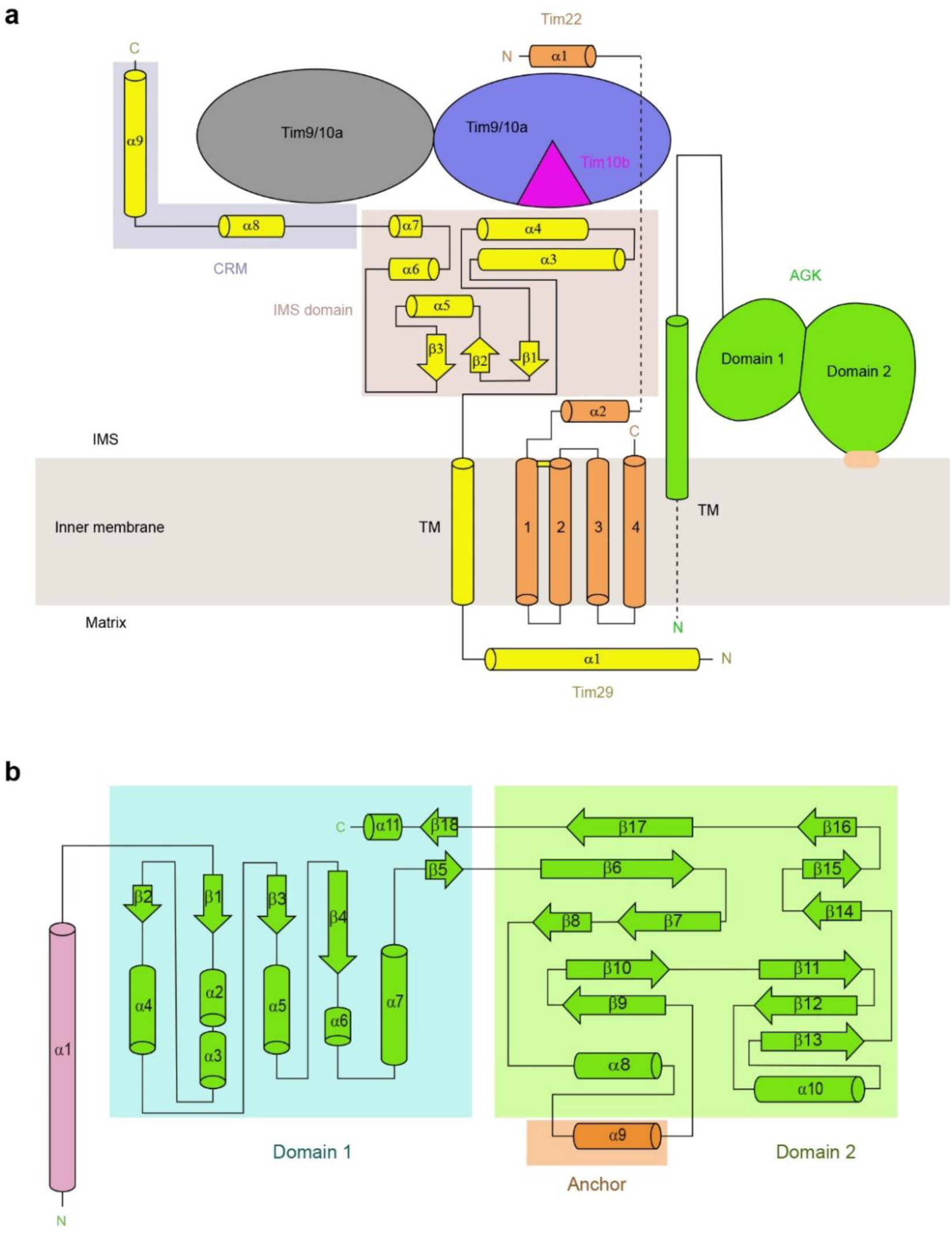
Topology diagrams of the TIM22 complex. **a**, Secondary elements are labeled. Tim29, yellow; Tim22, orange; AGK, chartreuse; Tim9/10a, slate; Tim9/10a/10b, blue; Tim10b, magenta. **b**, The topology diagram of AGK. α 1, light pink; anchor, orange; domain 1 (α2∼ α7, α12 and β1∼ β4) and domain 2 (α8∼ α10 and β5∼ β18).

**Extended Data Fig. 6.**
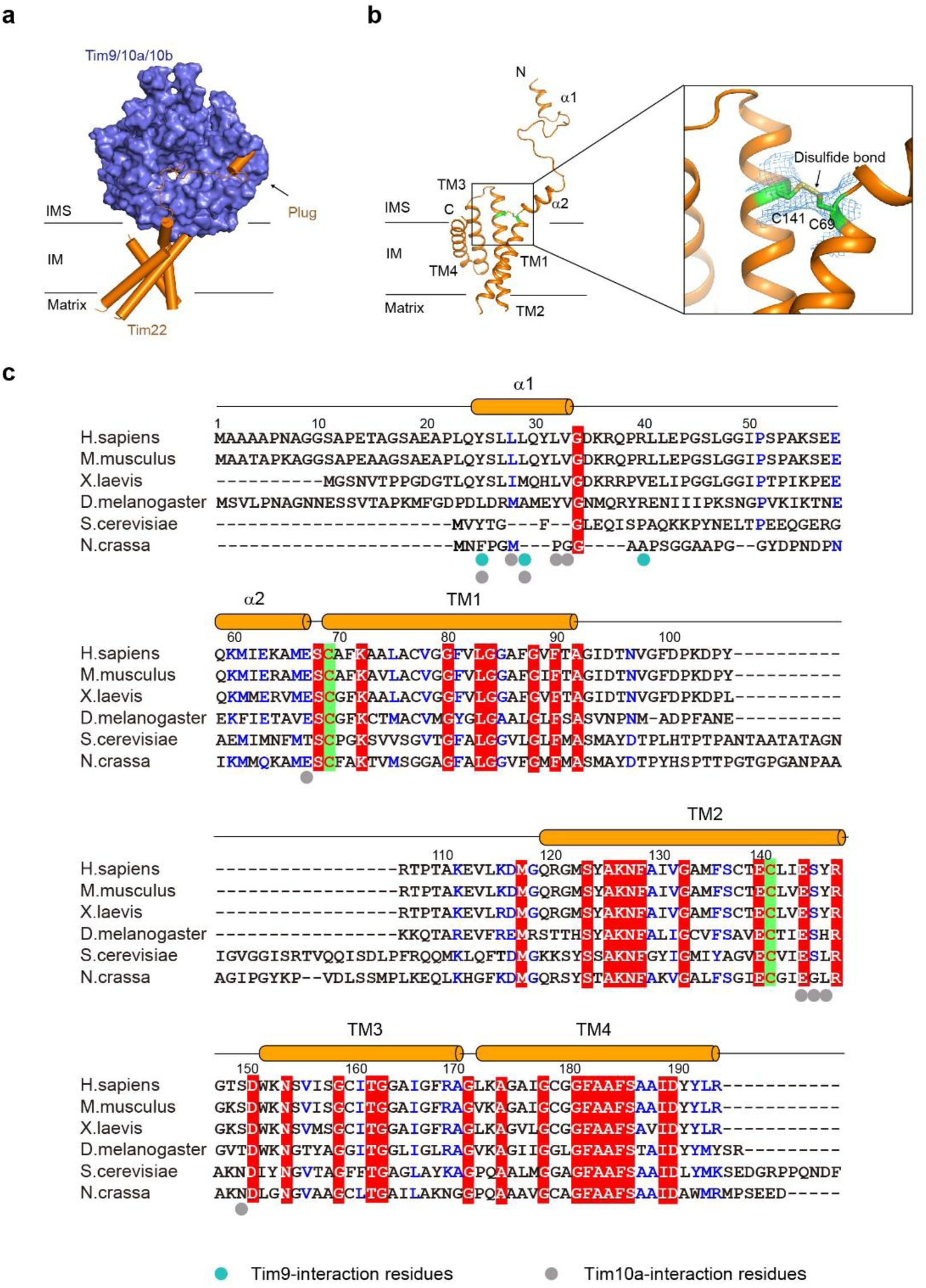
The N-terminal helix and disulfide bond in Tim22. **a**, Cartoon and surface model of Tim22 and Tim9/10a/10b. Plug indicates the first α-helix of Tim22. **b**, Disulfide bond between C69 and C141 of Tim22. **c**, The sequences (Uniprot code: *Homo sapiens*, Q9Y584; *Mus musculus*, Q9CQ85; *Xenopus laevis*, Q5U4U5; *Drosophila melanogaster*, Q8IN78; *Saccharomyces cerevisiae*, Q12328; *Neurospora crassa*, Q9C1E8) are aligned using ClustalW. Residues involved in the interaction with Tim9 and Tim10a are highlighted with cyan and slate balls. Cysteines forming disulfide bond are highlighted using yellow boxes. The secondary structural elements are indicated above the sequences.

**Extended Data Fig. 7.**
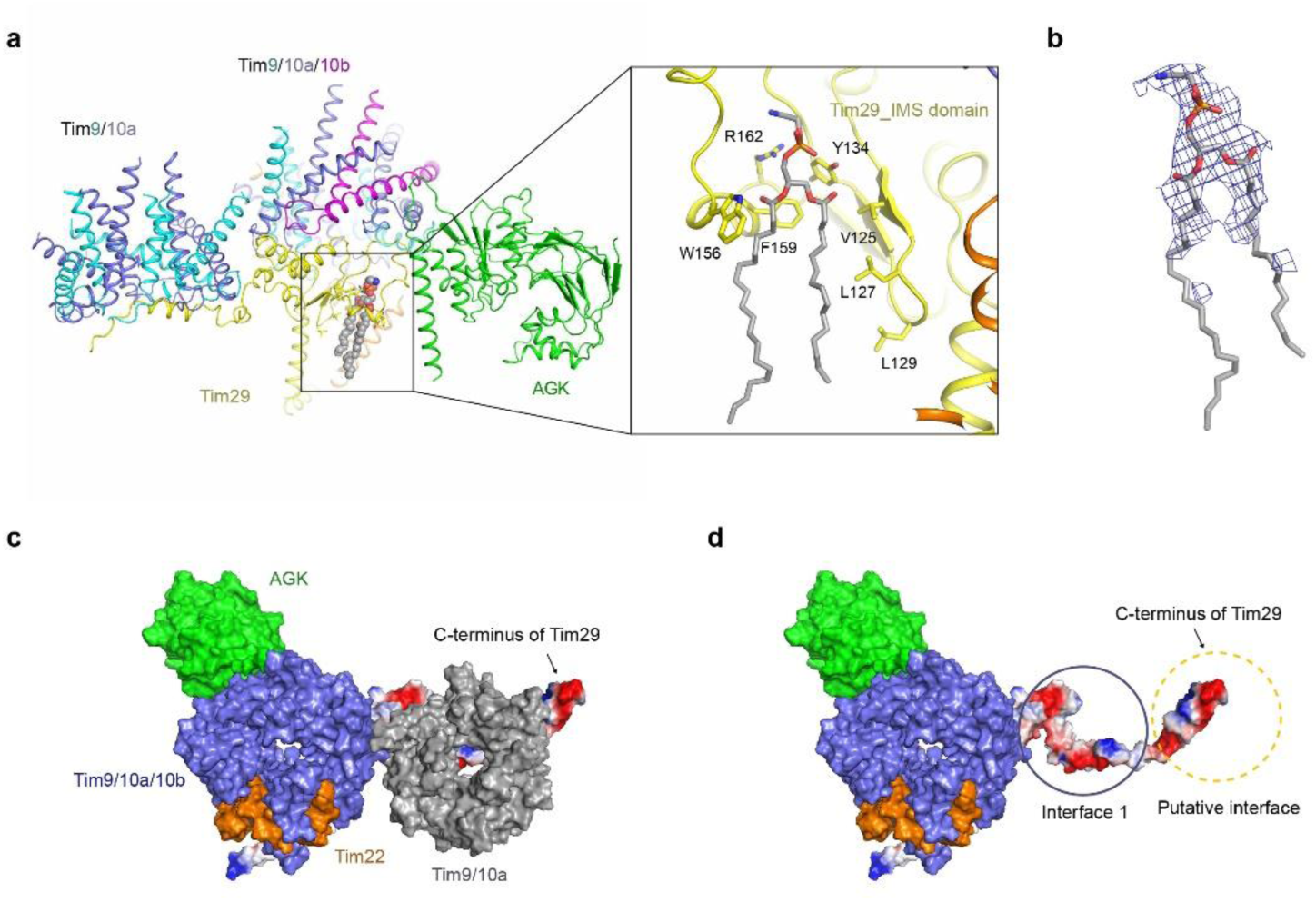
A phospholipid in the TIM22 complex and a new putative chaperone recruiting interface of Tim29. **a**, A phospholipid, binding at the interface between Tim29 and inner membrane. The aliphatic tails of the latter phospholipid may interact with several hydrophobic residues (V125, L127, L129, W156 and F159). The phosphate group likely to interact with several hydrophilic residues (Y134, and R162). **b**, The electron microscopy maps for the phospholipid resembling to the structure of Phosphatidylethanolamine (PDB code: PTY). **c, d**, Electrostatic surface of TIM29. A top view (from IMS to matrix) of TIM22 complex is shown as colored surface or electrostatic surface potential. The interface 1 is framed out using a slate circle. A putative interface for recruiting another chaperone is framed out using a yellow dashed circle. Tim9/10a is omitted for clarity in (D). AGK chartreuse; Tim22, orange; Tim9/10a/10b, blue; Tim9/10a, slate; Tim29, electrostatic surface potential.

**Extended Data Fig. 8.**
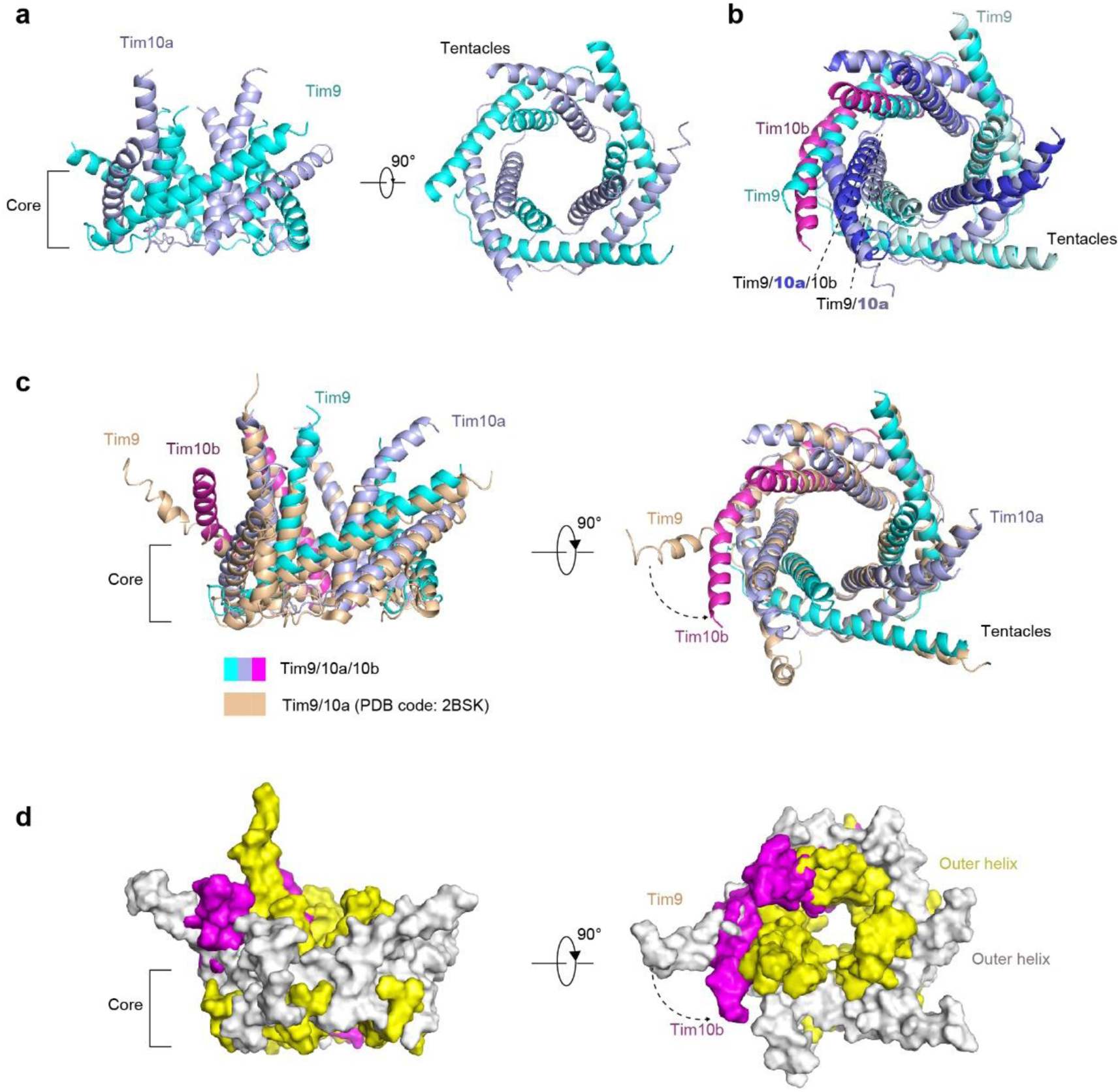
Center pore comparison of Tim9/10a and Tim9/10a/10b hexamer. **a**, Two perpendicular views of Tim9/10a/10b hexamer. **b**, Structural alignment of Tim9/10 and Tim9/10a/10b. Tim9, cyan; Tim10a, slate or peal blue; Tim10b, magenta. **c**, A structure alignment of chaperone Tim9/10a/10b with free chaperone Tim9/10a (PDB code: 2BSK). Tim9, cyan; Tim10a, blue; Tim10b, magenta; free chaperone Tim9/10a, wheat. **d**, A structure alignment of chaperone Tim9/10a/10b with free chaperone Tim9/10a (PDB code: 2BSK) are shown as colored surface: inner helices of Tim9/10a/10b, yellow; outer helices of Tim9/10a/10b and free Tim9/10a, slate; Tim10b, magenta.

**Extended Data Fig. 9.**
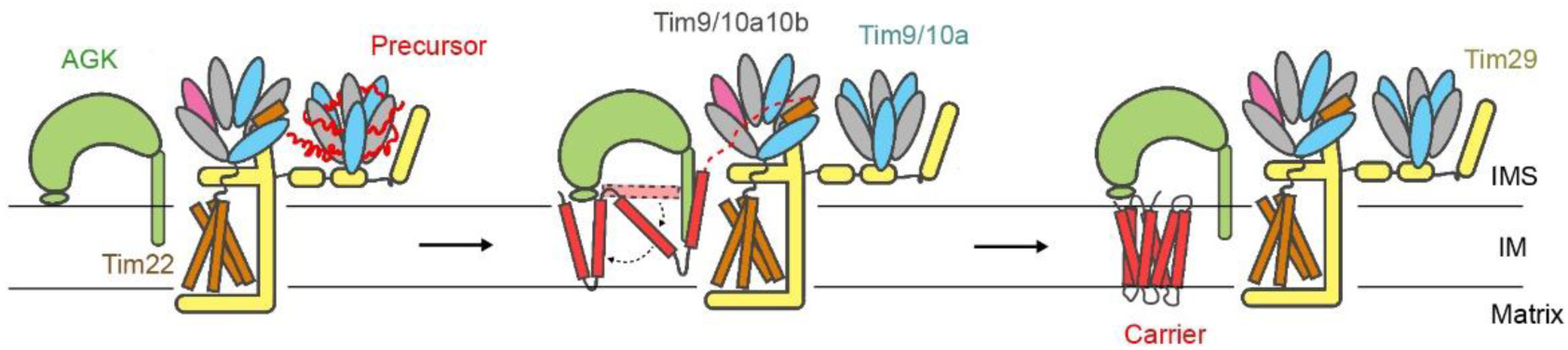
Implication on carrier precursor insert into inner membrane. Cartoon model of carrier precursor insertion into the membrane through The TIM22 complex. The carrier precursor (red line, left panel) wraps around the Tim9/10a chaperone and then inserts into the membrane (red stick, middle and right panel) through the cavity of AGK by a “bending-in” mechanism.

**Extended data Table 1.**
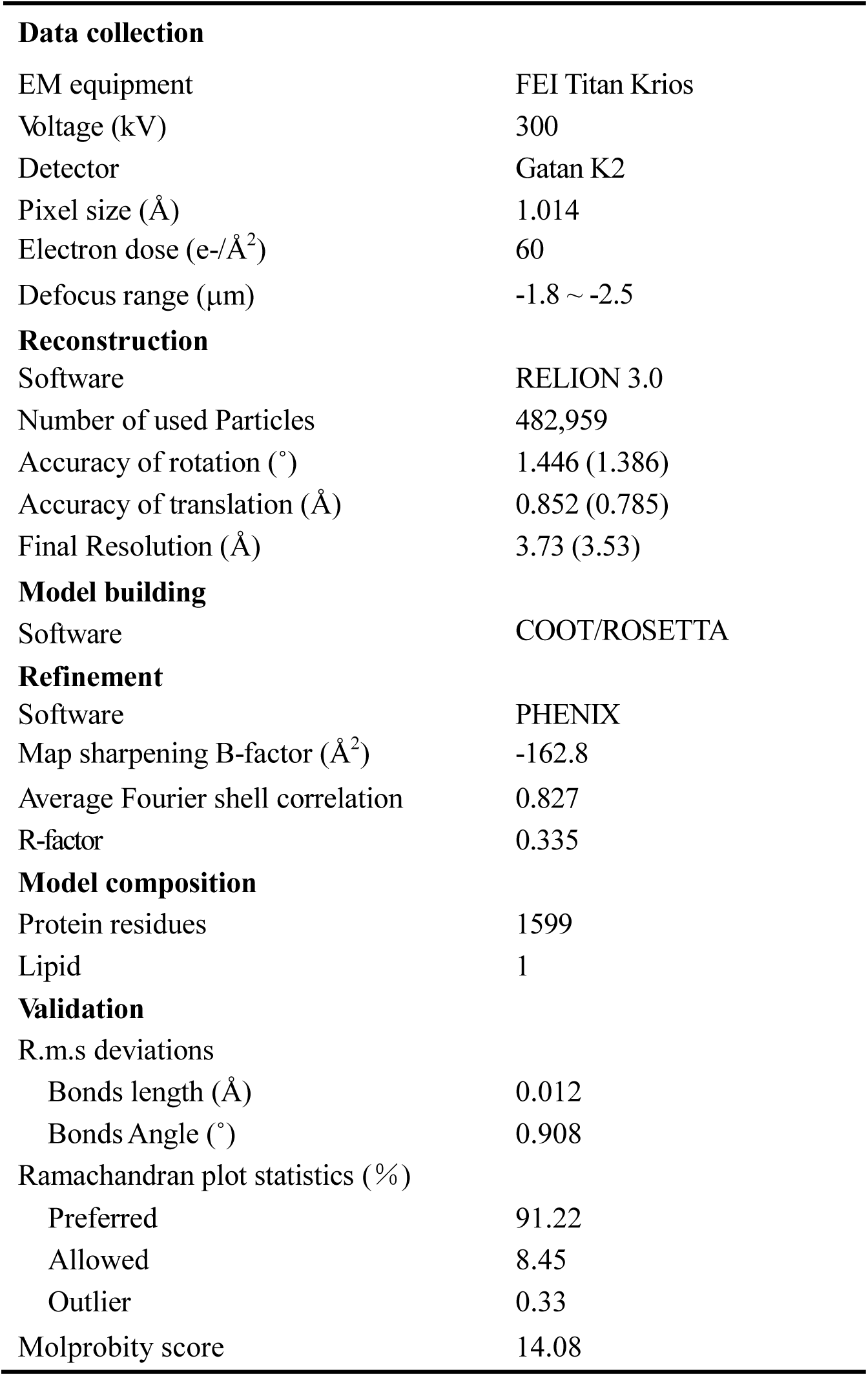
Statistics of Cryo-EM data collection and refinement of the human TIM22 complex.

**Extended data Table 2.** Summary of model building for the human TIM22 complex.

|  | Length | Domain | Copies | PDB code | Modeling<sup>1</sup> | Resolution (Å) | Chain ID |
| --- | --- | --- | --- | --- | --- | --- | --- |
| **Tim22** | 194 | 24-194 | 1 |  | De novo | 3.5-6.0 | A |
| **AGK** | 422 | 23-415 | 1 | 2QV7 | HM | 3.5-4.2 | B |
| **Tim29** | 260 | 21-244 | 1 |  | De novo | 3.2-5.2 | C |
| **Tim9** | 89 | 5-82 | 5 | 2BSK | RD | 3.2-4.0 | D-F, K, L |
| **Tim10a** | 90 | 2-80 | 6 | 2BSK | RD | 3.2-3.8 | G-I, M-O |
| **Tim10b** | 103 | 3-85 | 1 | 2BSK | HM | 3.2-3.8 | J |

<sup>1</sup>Abbreviations in the “Modeling” column are as follows. HM, homology modeling; RD, rigid docking and refinement; De novo, de novo model building.

